# Assessment of kinship detection using RNA-seq data

**DOI:** 10.1101/546937

**Authors:** Natalia Blay, Eduard Casas, Iván Galván-Femenía, Jan Graffelman, Rafael de Cid, Tanya Vavouri

## Abstract

Analysis of RNA sequencing (RNA-seq) data from related individuals is widely used in clinical and molecular genetics studies. Sample labelling mistakes are estimated to affect more than 4% of published samples. Therefore, as a method of data quality control, a way to reconstruct pedigrees from RNA-seq data would be useful for confirming the expected relationships. Currently, reconstruction of pedigrees is based mainly on SNPs or microsatellites, obtained from genotyping arrays, whole genome sequencing and whole exome sequencing. Potential problems with using RNA-seq data for kinship detection are the low proportion of the genome that it covers, the highly skewed coverage of exons of different genes depending on expression level and allele-specific expression.

In this study we assess the use of RNA-seq data to detect kinship between individuals, through pairwise identity-by-descent (IBD) estimates. First, we obtained high quality SNPs after successive filters to minimize the effects due to allelic imbalance as well as errors in sequencing, mapping and genotyping. Then, we used these SNPs to calculate pairwise IBD estimates. By analysing both real and simulated RNA-seq data we show that it is possible to identify up to second degree relationships using RNA-seq data of even low to moderate sequencing depth.

## Introduction

Gene expression data from related individuals is used in genetics for a variety of purposes such as to test the heritability of gene expression traits (1) or to study the mechanisms of epigenetic inheritance of phenotypic traits (2) among others. Nowadays, gene expression is usually quantified by high throughput sequencing (RNA-seq). In many of these studies, multiple RNA samples extracted from different individuals from multiple families are processed in parallel. During sample and data processing, sample mislabelling can occur due to human error. This is detrimental for downstream analysis and has been estimated to affect at least 4% of published samples (3, 4). Given the importance of this problem, there are numerous programs and methodologies that deal with sample mislabelling by comparing paired samples from the same individual (5, 6) or using a heuristic data perturbation strategy (4). However, so far it is unclear whether kinship can be predicted directly from RNA-seq data so as to use kinship detection as a sample quality control step during data processing.

There are several methods to predict or confirm familial relationships and build heritability estimates from genetic data. Although these methods were initially based on microsatellites (7) and amplified fragment length polymorphism (AFLPs) (8), nowadays they usually rely on SNPs. Among the different methodologies for kinship detection, we can find methods that exclude impossible parent-offspring relationships (9–11), methods that calculate kinship coefficients and/or identical-by-descent (IBD) probabilities (12, 13) and likelihood methods (14, 15). RNA-seq data can be used to identify single nucleotide polymorphisms (16). Yet, the use of RNA-seq data for kinship detection and pedigree reconstruction has not been properly assessed so far. Some of the concerns with using this type of data for kinship detection are the limited number of SNPs adequately covered by reads and allelic imbalance (17, 18) that may hinder genotype calling by masking one of the two alleles at many variant positions along the genome.

Here, we report the results of our assessment of the utility of human RNA-seq data for kinship detection and pedigree reconstruction. Our input data is a set of raw RNA-seq data files and a set of known common variant positions. First, we filter both the RNA-seq data and the known variants to obtain reliable predicted SNPs for each RNA sample. We then use these SNPs to detect and represent familial relationships by estimating pairwise probabilities of identity-by-descent (IBD) (12, 19). By analysing human empirical as well as simulated RNA-seq data we show that the estimated IBD probabilities allow kinship detection and pedigree reconstruction, detecting up to second degree relationships even with low sequencing depth.

## Materials and Methods

### Retrieval of empirical data

We used three types of empirical data (**Figure 1**). First, we used previously published RNA-seq data from Epstein-Barr-virus-transformed peripheral blood B lymphocytes of human CEPH/UTAH family 1463 (20). This family has 17 individuals from three different generations, four grandparents, two parents and eleven children. Second, we retrieved genetic variation data from six pairs of first and second degree *relatives* from the CDX population of the 1000 Genomes Project. From this data we then simulated RNA-seq reads (the method of simulation of RNA-seq reads is described below). Third, we retrieved genetic variation data from unrelated individuals from the 1000 Genomes Project (21) from which we simulated families (the method of simulation of genotypes of family members is described below) and RNA-seq reads.

**Figure 1:**
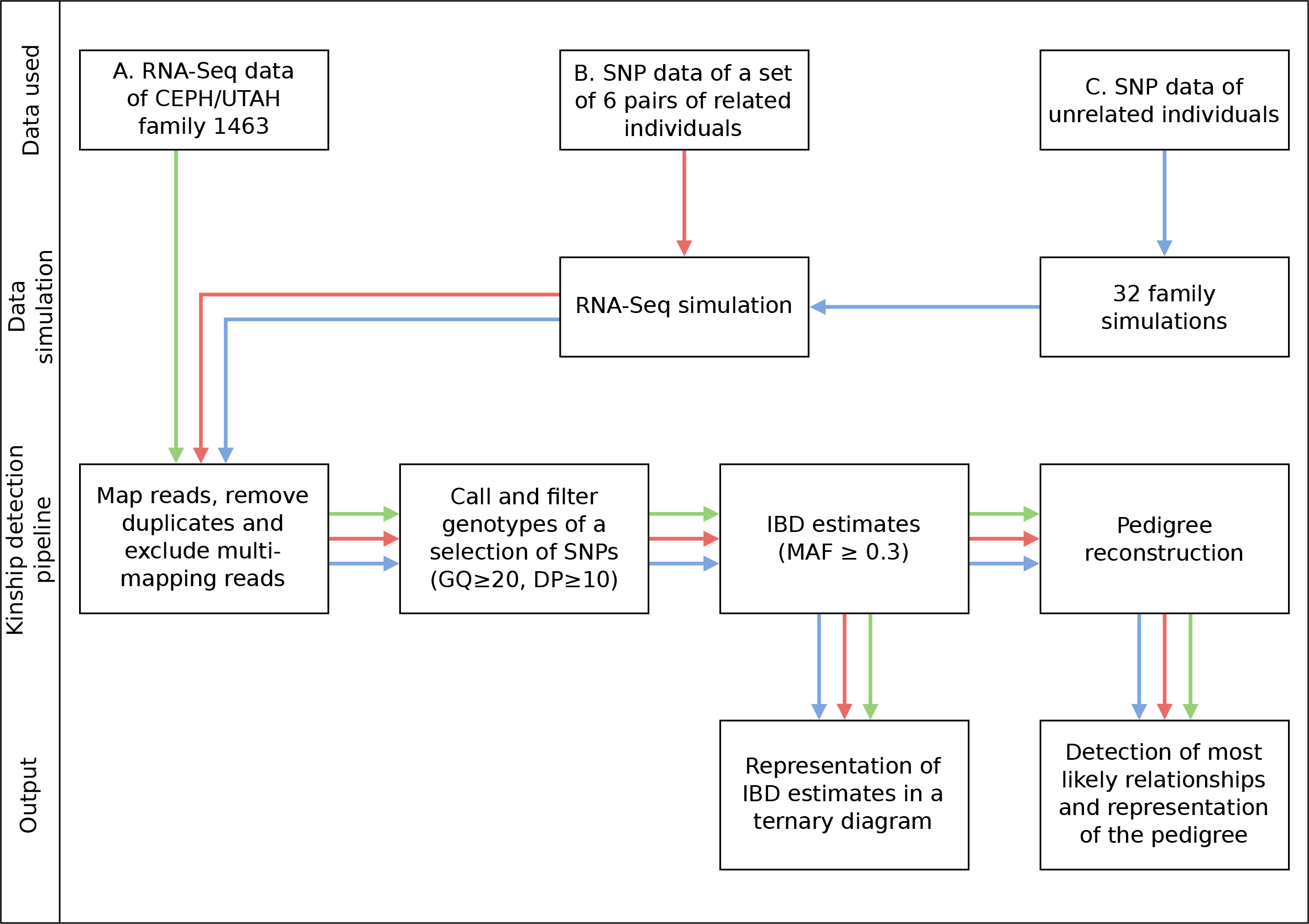
Overview of data workflow for kinship detection using RNA-seq data. GQ: genotype quality, DP:depth, IBD: identity-by-descent, MAF: minor allele frequency.

### Simulation of genotypes of family members

To simulate the genotypes of family members we used SNP data from the 1000 Genomes Project (21). We selected unrelated individuals as founders and used their haplotype data to simulate the rest of the family. We simulated data according to different pedigrees in order to assess the effect of pedigree complexity. Family simulations were carried out following Mendelian laws and taking into account linkage disequilibrium (assuming 1cM per Mb) with a custom R script.

### Simulation of RNA-seq reads

To simulate gene expression data from real or simulated individuals, we generated paired-end RNA-seq reads with flux-simulator v1.2.1 (22). We used two genome fasta files per individual (one per haplotype, including the SNPs to the reference genome with GATK v3.8 FastaAlternateReferenceMaker (23)) and the expression profile of B lymphocytes (custom .pro file with RPKMx150 obtained from the founders of CEPH/UTAH family 1463 (20) to get a sequencing depth of 40M reads per individual). We ran the simulation with library preparation and sequencing steps (-ls options) for the first haplotype. For the second haplotype we used the library (.lib file) of the first haplotype and ran only the sequencing step (-s option) to make sure the same genes are expressed in both haplotypes.

### Data filtering, read mapping and kinship detection

We aligned empirical and simulated RNA-seq reads to the reference genome hg19 using HISAT2 v2.1.0 (24), removed duplicates and kept only uniquely mapping reads. We retrieved 14.8 million common SNPs from the UCSC Genome Browser (dbSNP Build ID 150)(25). We used BEDtools intersect v2.26.0 (26) to identify and remove 38,572 SNPs in 83 imprinted genes (http://www.geneimprint.org/ accessed: Mar 8, 2018)(27) and 8.2 M SNPs in repeats (RepeatMasker annotation downloaded from the UCSC Genome Browser), ending up with 6.2 M SNPs. Genotypes were obtained with SAMtools mpileup v1.7 and BCFtools call v1.4 (28). We used VCFtools v0.1.14 (29) to select only those SNPs with a depth ≥ 10 (option --minDP 10) and a genotype quality ≥ 20 (option --minGQ 20). To determine pairwise IBD estimates we used PLINK v1.9 (30), considering only autosomal SNPs with a minor allele frequency ≥ 0.3 (--maf 0.3 option) to obtain an optimal separation between groups. Representation of pairwise IBD estimates was done in R v3.5.0 (31) with the method described by Galván-Femenía et al. (2017) (32). Using the IBD estimates obtained in the previous step, we used PRIMUS v1.9.0 (33) to predict pairwise relationships and to reconstruct the whole pedigree, considering as unrelated those pairs of individuals with a coefficient of relatedness lower than 0.2 (option −t 0.2). Sex data was also provided. The sex of the individuals from the real data was inferred from counting reads mapping on the Y chromosome and compared against the reported sex.

## Results

### Analysis workflow for kinship detection using RNA-seq from related individuals

To assess how feasible it is to detect kinship using RNA-seq data we used three types of data (**Figure 1 and 2**). First, we used a previously published set of RNA-seq data from B-lymphocytes from members of an extended human family (**Figure 2A**). Second, we used available SNP data from six pairs of first and second degree relatives from the CDX population of the 1000 Genomes Project from which we simulated RNA-seq data (**Figure 2B**). Third, we used available SNP data from unrelated individuals from which we simulated the genotypes of family members and the corresponding RNA-seq samples (**Figure 2C**). We simulated four types of pedigrees to assess the effect of pedigree complexity. After data retrieval or simulation, we used the same analysis workflow which in brief consisted of mapping and filtering RNA-seq reads, calling and filtering genotypes of a set of filtered SNPs, calculating IBD estimates and finally visualising the IBD estimates in ternary diagrams or providing the IBD estimates to the software PRIMUS for prediction of pairwise relationships and pedigree reconstruction.

**Figure 2:**
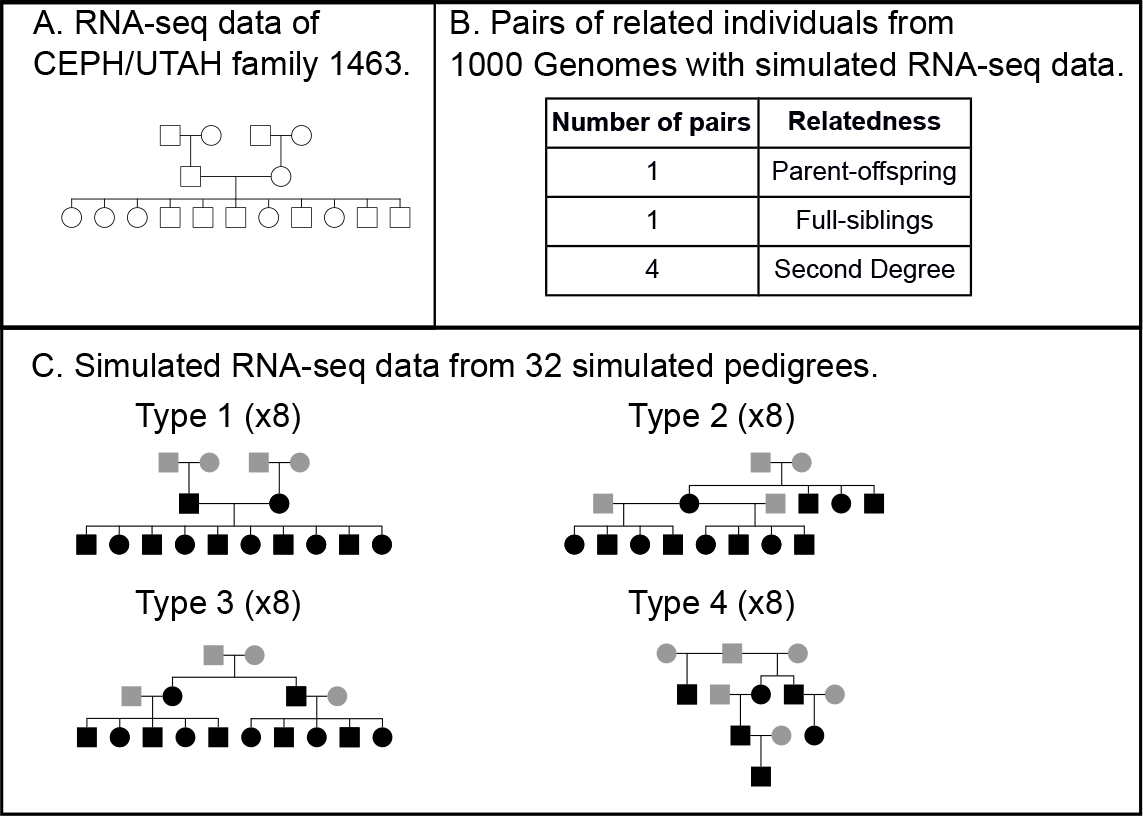
Structure of datasets used for the assessment of kinship detection using RNA-seq data. (A) Structure of CEPH/UTAH family 1463 with empirical RNA-seq data. (B) List of real related pairs of individuals with simulated RNA-seq data. (C) Simulated families with simulated RNA-seq data (pedigree types 1 - 4). Real individuals with simulated RNA-seq data are highlighted in grey. Simulated individuals with simulated RNA-seq data are highlighted in black.

### Kinship detection using empirical RNA-seq data from a human family

We mapped and filtered the raw RNA-seq data from B-lymphocytes from members of CEPH/UTAH family 1463 and in parallel filtered the initial set of known common human variants as described in the methods. Reads were mapped to the reference human genome. This step excluded on average 12% of the initial reads (range 6-24%, depending on the individual) that could not be mapped. We then removed PCR duplicates as they can potentially amplify sequencing errors. This step led to the removal of an average of 21% of the reads (range 17-30%). We also removed multi-mapping reads to avoid mapping errors, excluding on average 4.6% of the reads (range 4.1-5.3%). We filtered the known common human variants; we removed SNPs in repeats and in imprinted genes (55% and 0.26% of SNPs excluded, respectively) that could bias genotyping. Having obtained a filtered set of RNA-seq reads (a mean of 34M reads per individual) and SNPs (6.2 M SNPs), we proceeded to call and filter genotypes, obtaining 633,636 SNPs that were covered by reads in at least one individual. To exclude those SNPs with low coverage we selected a minimum depth of 10 reads per SNP, and to avoid genotyping errors we filtered out those SNPs with a genotype quality lower than 20, which means an error rate of 0.01 (in this step we excluded 76% of the SNPs). Finally, we considered those SNPs with a minor allele frequency of at least 0.3 (excluding 89.5% of the SNPs). We ended the filtering process with 16,004 SNPs present in at least one individual. The number of SNPs that were used for downstream analyses was between 5,657 and 8,600, depending on the number of missing genotypes observed for each pair of individuals.

We used ternary diagrams to visualize the estimated probabilities of sharing 0, 1 and 2 IBD alleles (referred here as Z0, Z1 and Z2 respectively) meaning that none, only one or both alleles of the pair are inherited from the same recent common ancestor. In ternary diagrams, the coordinates of each data point are the three IBD estimates for each pair of individuals in the dataset. We observed that the different relationships were grouped around their theoretical values with no overlap between them. As expected, parent-offspring pairs are clustered at the Z1 corner (meaning that they share 1 IBD allele at all SNPs), unrelated individuals at the Z0 corner (they do not share any IBD alleles), second degree relationships on the Z0-Z1 axis (they share 1 IBD allele at half of their SNPs) and full-siblings in the middle of the graph (they can share from 0 to 2 IBD alleles per SNP) (**Figure 3A, Supplementary Table 1**).

**Figure 3:**
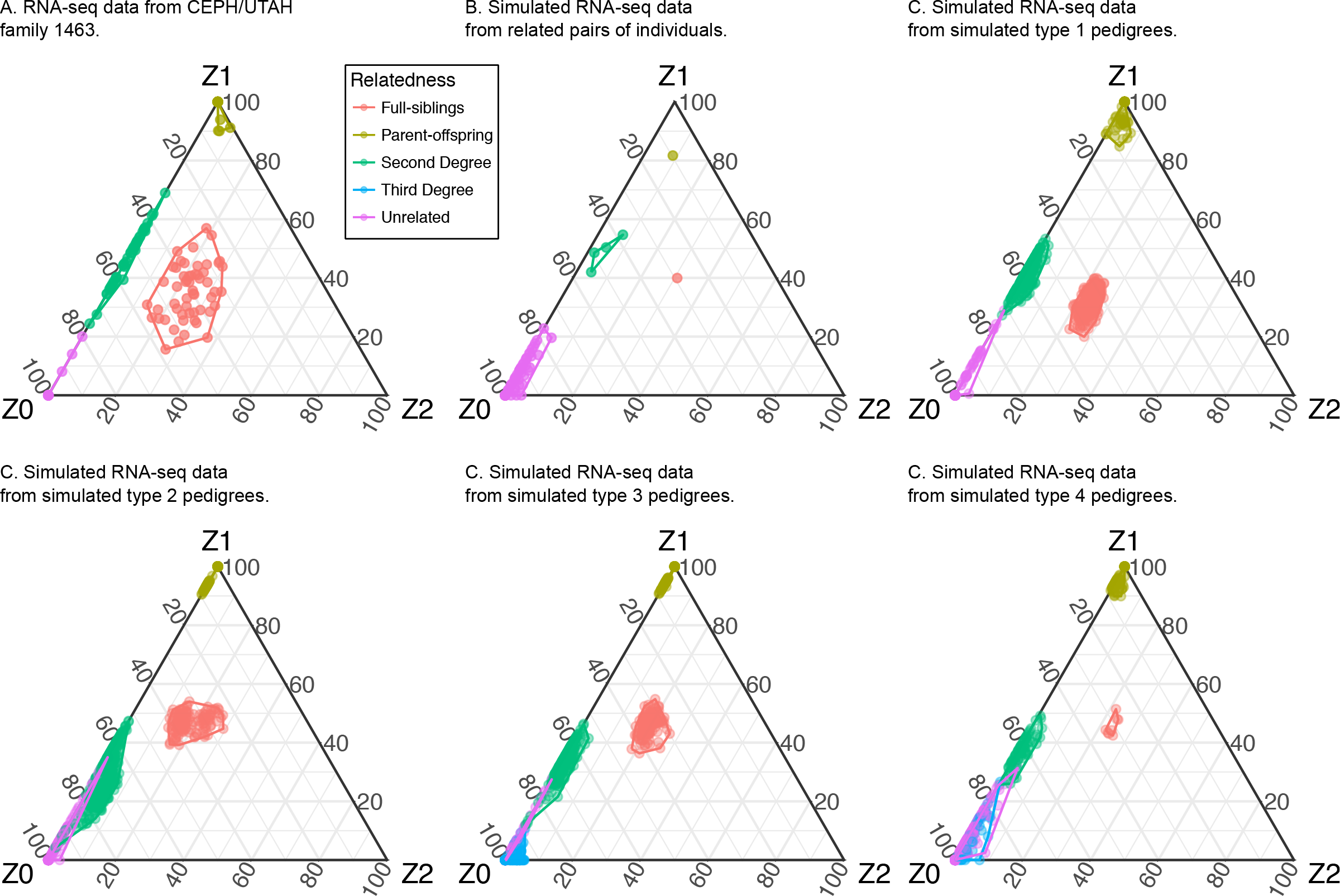
Ternary diagrams of IBD estimates for (A) CEPH/UTAH family 1463, (B) simulated RNA-seq data from real related pairs of individuals, and (C) simulated RNA-seq data from simulated related individuals from different pedigree types (type 1 - type 4).

The actual pedigree was also correctly reconstructed with PRIMUS (33) from the estimated IBD values. In addition to the top scoring pedigree, which in this case corresponds to the correct one, PRIMUS also reports the predicted pairwise relationships. In this case it did not correctly identify some of the pairwise relationships; the mis-identified pairs were 2% of full sibling pairs (one pair was classified as second degree relatives), 32% of second degree pairs and 9% of unrelated pairs of individuals.

According to these results, we can conclude that kinship detection is possible with RNA-seq data at least for this dataset, allowing us to detect and represent first and second degree relationships. Although the sensitivity decreases as the degree of relatedness increases, pedigree reconstruction is robust.

### Impact of sequencing depth on kinship detection

The sequencing depth of the real RNA-seq data from the extended human family ranged from 37 to 67 M reads, with a mean depth of 52 M. To understand the effect of sequencing depth on kinship detection, we randomly subsampled different numbers of reads from the FASTQ input files. We subsampled 37 M reads (the number of reads for the individual with the lowest coverage) to homogenize the sequencing depth for all individuals and then also 30 M, 20 M, 10 M, 8 M, 6 M, 4 M and 2 M reads from each sample to determine the minimum sequencing depth for kinship detection and pedigree reconstruction.

Visual inspection of estimated probabilities using ternary diagrams (**Figure 4**) revealed that there was no overlap between second degree relatives and unrelated individuals when using RNA-seq data with 52, 37, 30, 20, 10 and 6 M reads. In the simulation of 8 M reads we observed a slight overlap between second degree relatives and unrelated individuals. In the simulation of 4 M reads, there was overlap between second degree and unrelated pairs of individuals and also between second degree and full siblings. With 2 M reads all groups overlapped except parent-offspring and unrelated pairs of individuals.

**Figure 4:**
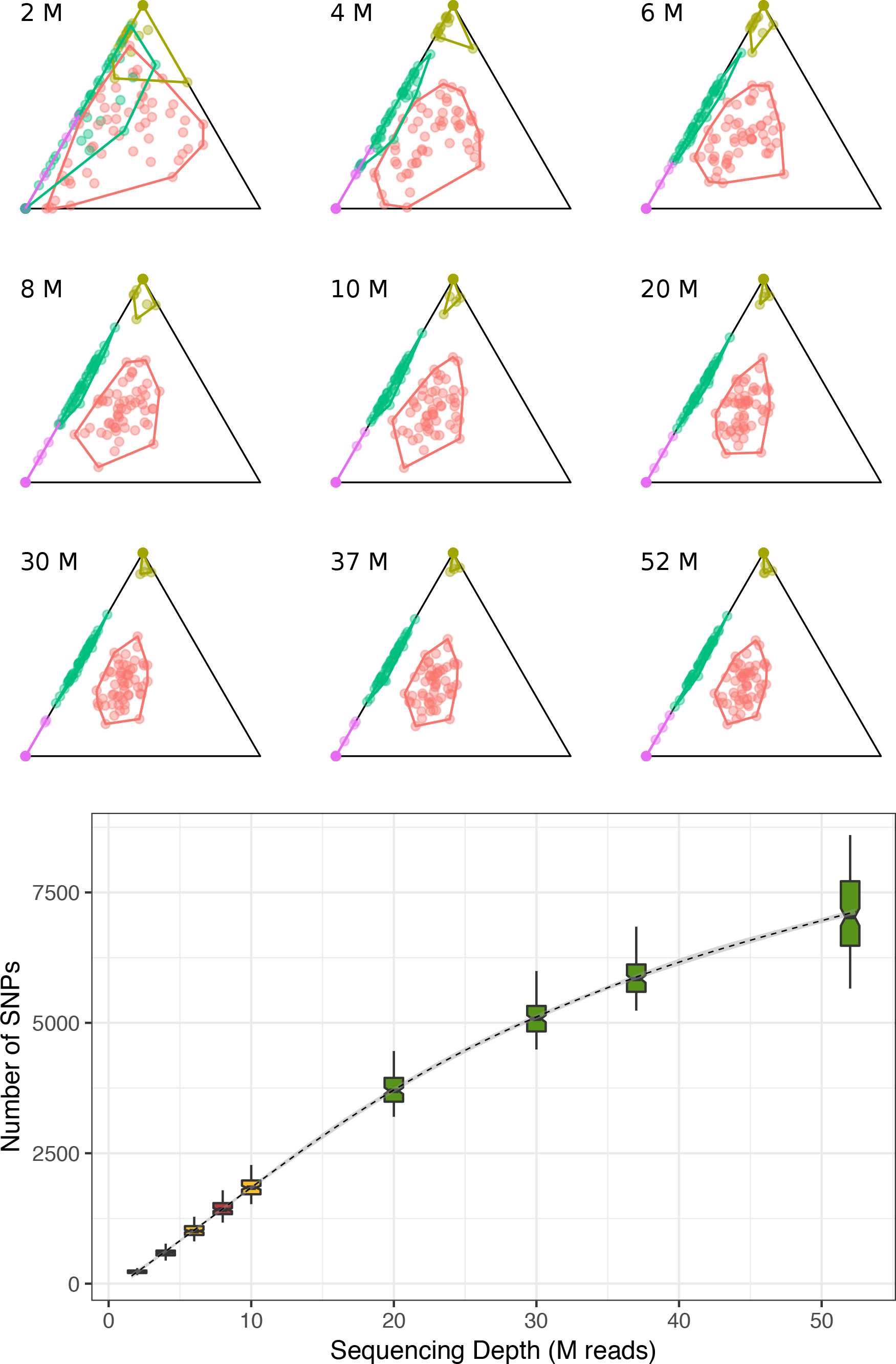
Ternary diagrams of IBD estimates for each sequencing depth (top) and number of SNPs used for pairwise comparison per sequencing depth (bottom). Green colour indicates that the pedigree was correctly reconstructed, yellow colour indicates that there was no overlapping between groups but the pedigree reconstruction was not possible, and red colour indicates that there was an overlapping between groups. The box with a sequencing depth of 52, is the one for the original data, being the mean for the different sequencing depths (from 37 to 67 M reads).

Pedigree reconstruction was only possible for RNA-seq data with 52, 37, 30 and 20 M reads because, when decreasing the sequencing depth, the probability to share one IBD allele (Z1) of some full siblings decreased way below the expected values (for example Z0=0.6741, Z1=0.0712, Z2=0.2547).

Taken together, these results demonstrate that sequencing depth affects the number of SNPs used for pairwise comparison, but overall its effect on kinship detection is small, providing acceptable IBD estimates (no overlapping between relationships) even with 6 M reads, where the number of SNPs is as low as 811-1,282 (**Figure 4**). As IBD estimates with 8 M reads were not as good, we conclude that the minimum sequencing depth to be able to detect kinship in RNA-seq data through IBD probabilities is 10 M reads. In contrast, the minimum sequencing depth to be able to reconstruct the correct pedigree is 20 M reads.

### Kinship detection using empirical genotypes of related individuals with simulated RNA-seq data

To further assess the feasibility of estimating relationships from RNA-seq samples from related individuals, we simulated RNA-seq data from real related individuals with known genotypes. We used variation data from six pairs of first and second degree relatives from the CDX population of the 1000 Genomes Project (**Figure 2B**). We simulated paired-end RNA-seq reads with the expression profile of B-lymphocytes. We obtained this gene expression profile from the mean of the number of reads mapped to a gene per gene length in kilobases per million mapped reads (RPKM) of the founders of the CEPH/UTAH family 1463. Then, we multiplied those values by a factor of 150 to simulate the desired expression profile with a sequencing depth of 40 M reads. As expected, the simulated RNA-seq data is highly correlated with the real data (**Supplementary Figure 1**) - pearson’s correlation estimates are on average 0.918 (range 0.913-0.925). We used the same workflow described for the empirical RNA-seq data. The resulting number of SNPs used for the pairwise comparisons was between 4,501 and 4,632.

In agreement with the results obtained from the real RNA-seq data, visual inspection of estimated probabilities in ternary diagrams revealed that all relationships are clearly separated (**Figure 3B**). Similarly, the pedigree reconstruction program was able to correctly classify and represent all relationships in a pedigree. We conclude that kinship detection using RNA-seq data is possible using different datasets; for entire families as well as for pairs of related individuals.

### Kinship detection using simulated genotypes of family members with simulated RNA-seq data

We then asked whether the complexity of the pedigree could affect kinship detection and pedigree reconstruction. To address this, we generated additional RNA-seq data from different types of pedigrees, this time simulating both gene expression and also the genotypes of individuals in the pedigrees. We used genotypes of real unrelated individuals from the 1000 Genomes Project as founders (**Figure 2C**, genotypes of real individuals shown in grey, simulated genotypes of offspring shown in black). We simulated the families by using data from eight different populations from the 1000 Genomes Project: GBR, KHV, IBS, LWK, CLM, CDX, PEL and ACB; and four pedigree structures including different degrees of relationship (**Figure 2C**):

- Type 1: A 16-member pedigree with the same relationships as CEPH/UTAH family 1463.
- Type 2: A 16-member pedigree with all first and second degree relationships from **Supplementary Table 1**.
- Type 3: A 16-member pedigree with first, second and third degree relationships.
- Type 4: A 12-member pedigree with all first, second and third degree relationships from **Supplementary Table 1**.

We made 32 simulations, one for each pedigree type and population. Sex was not simulated as sex chromosomes are not used for IBD estimation. After data filtering, we observed that the number of SNPs used for pairwise comparison was higher in African populations (ACB and LWK) ranging from 5,180 to 5,974 and lower in Asian populations (KHV and CDX) ranging from 4,289 to 5,084 **(Supplementary Figure 2**).

Visual inspection of IBD probabilities (**Figure 3C**) revealed that first degree relationships were clearly separated from other relationships, second degree relationships were separated from unrelated individuals in all simple pedigrees (type 1 simulated pedigrees) and some type 2, 3 and 4 simulated pedigrees. In contrast, third degree relationships overlapped unrelated pairs of individuals in all pedigrees where they were present (**Figure 3C**, pedigree type 3 - type 4).

Reconstruction of the actual pedigree was possible for all simulated type 1 pedigrees, seven type 2 pedigrees and four type 4 pedigree (one of them was not the top scoring one, but the fourth in the PRIMUS output), but for none of the type 3 pedigrees (**Figure 5A**). Although PRIMUS was not able to reconstruct all the simulated pedigrees, it was able to identify all first degree relatives, 35% of second degree relatives (in type 1 pedigrees this percentage was 68%), 12% of third degree relatives, and 93% of unrelated individuals (**Supplementary Figure 3**).

**Figure 5:**
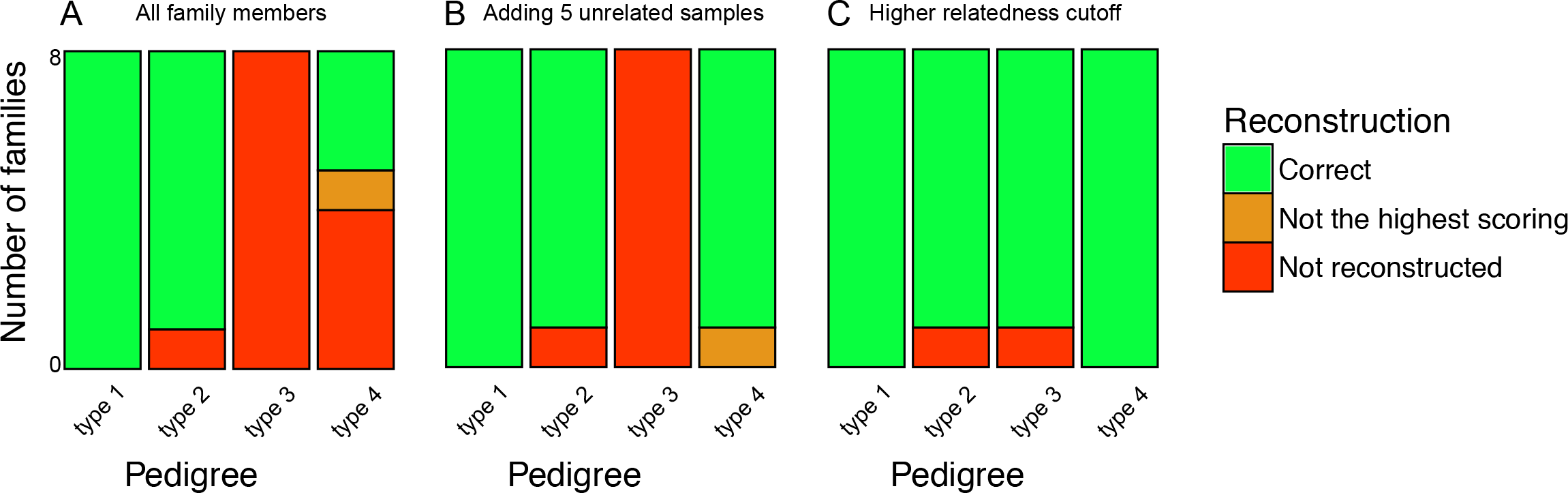
Number of simulated pedigrees reconstructed when analyzing (A) the original samples with our default workflow (B) the original samples with five additional unrelated individuals (C) and when analyzing the original samples with a higher relatedness cutoff (0.375 instead of 0.2).

We conclude that family structure affects kinship detection, providing good IBD estimates for simple families with up to second degree relatives, but in pedigrees with third degree relatives there is overlap between different relationships. Accordingly, entire pedigree reconstruction works better for simple families, getting worse when the number of third degree related pairs increases (type 3 pedigree has 25 pairs of third degree related individuals while type 4 pedigree has 6). Population specific differences in genetic variability lead to different number of SNPs that pass all filters, however these differences do not affect the results.

We wondered whether the addition of unrelated individuals would improve kinship detection. Presumably, they include more variability and make related individuals look more similar to each other. To test this, we added 5 unrelated individuals from the same population to each simulated family. The result was just as good for all families that were previously correctly reconstructed but it improved for four families, reconstructing correctly all type 1, seven type 2, all type 4 pedigrees but none of type 3 (**Figure 5B**). We conclude that adding data from unrelated individuals would be useful in those cases where the reconstruction is not possible.

Last, we asked whether it was possible to correctly predict the pedigree structure using only the most related individuals in complex pedigrees by using a stricter relatedness cutoff. We reran PRIMUS with a higher relatedness cutoff (option −t 0.375, instead of the previously used −t 0.2) and found that type 1 and type 2 pedigrees were reconstructed equally well but also that 7 out of 8 type 3 pedigrees and all type 4 pedigrees could also be correctly reconstructed (**Figure 5C**). We conclude that a more restrictive relatedness cutoff works better than adding unrelated individuals in those cases where most or all of the individuals of the pedigree are connected through a first degree relative.

## Discussion

We have shown here that kinship detection based on estimates of identity-by-descent probabilities using RNA-seq data from B-lymphocytes is possible allowing the detection of up to second degree relationships. In addition, we have shown that the actual pedigrees can be fully reconstructed and, although pedigree reconstruction works better for simple pedigrees, some pedigrees with third degree relatives can also be reconstructed. Furthermore, using simulations, we have shown that the ability to detect kinship between individuals is not limited by RNA-seq sequencing depth since it is still possible at a depth significantly below what is considered acceptable for gene expression analyses (usually 50 M reads). Reconstruction of full pedigrees requires higher sequencing depth but is still possible with at least 20 M reads.

There are additional issues that remain to be investigated in future studies. One of them is whether data from other tissues or cell types would result in better or worse kinship detection. Theoretically, RNA samples with moderate expression of many genes would lead to a better performance than those with high expression of few genes (as expected in highly specialized differentiated cell types), as the genomic coverage would be higher in the former. Another issue is whether gene expression data from other species would provide similar results to the ones from humans. Genetic variability among different individuals from a particular species or a particular population could play an important role in kinship detection with higher genetic diversity between individuals leading to better results.

In conclusion, we have shown here that RNA-seq data can be used to identify samples from closely related individuals using estimated identity-by-descent probabilities calculated from predicted SNPs. It turns out that the effect of calling genotypes correctly at the single nucleotide level due to allele-specific expression is not large enough to impede kinship detection from IBD estimates. We therefore recommend the estimation of IBD probabilities and visualization of the clustering of samples in ternary graphs or the use of pedigree reconstruction programs such as PRIMUS as a quality control step in studies that generate multiple RNA-seq samples from related individuals.

## Acknowledgements

We thank the IGTP High Performance Computing Platform (IGTP HPC) for computing resources and Iñaki M de Ilarduya and Lloyd Goodman for technical support.

