## Supplementary Table and Figures for "Assessment of kinship detection using RNA-seq data"

**This PDF file includes:**

Supplementary Table 1

Supplementary Figure 1

Supplementary Figure 2

**Supplementary Table 1.** Expected IBD probabilities of different relationships and degrees of relatedness. Z0, Z1 and Z2 represent the probabilities that a pair of individuals share 0, 1 or 2 identical by descent alleles respectively.

| Degree | Relationship | Z0 | Z1 | Z2 |
|---|---|---|---|---|
| 1 | parent-offspring | 0 | 1 | 0 |
| | full-siblings | 0.25 | 0.5 | 0.25 |
| 2 | grandparental, avuncular, half-siblings | 0.5 | 0.5 | 0 |
| 3 | First-cousins, great-grandparental, great-avuncular, half-avuncular | 0.75 | 0.25 | 0 |
| ∞ | unrelated | 1 | 0 | 0 |

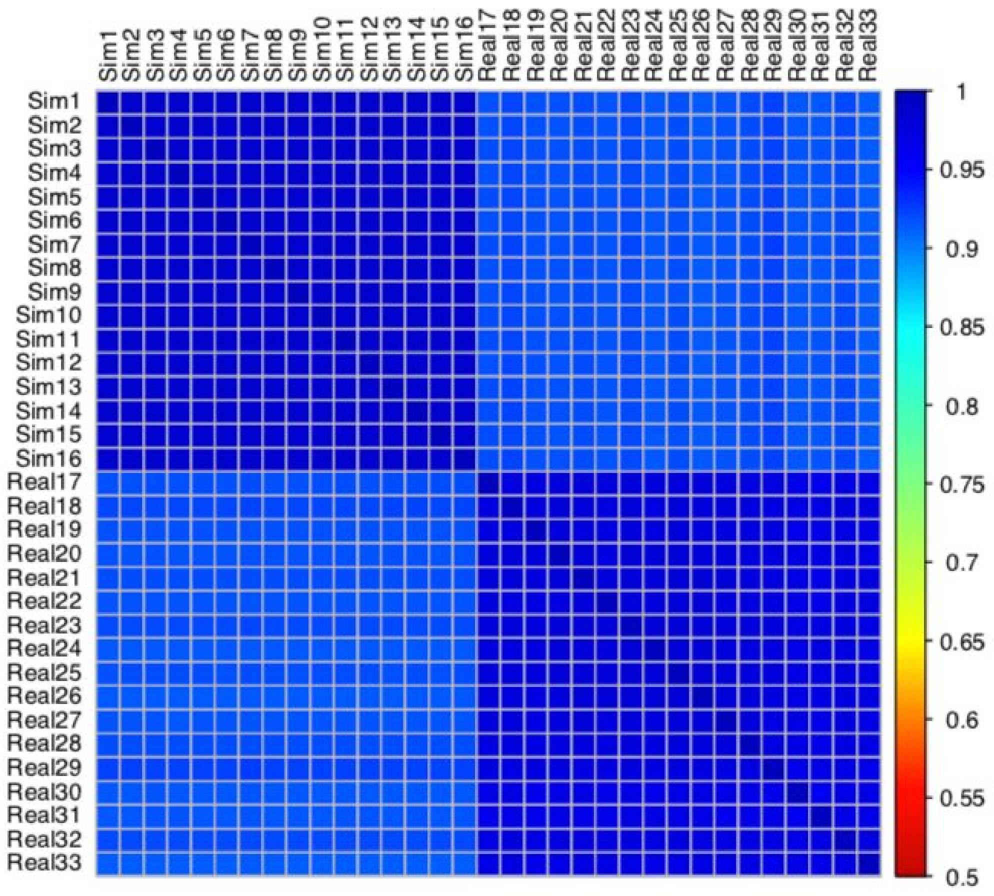


**Supplementary Figure 1.** Correlation matrix of the number of reads per gene in logarithmic scale between Simulated data (Sim1 - Sim16: individuals 1 - 16 of a type 1 simulated family) and Real data (Real1 - Real17: SRR1258217 - SRR1258233). Mean correlation between Real data: 0.975, between Simulated data: 0.988, between Real data and Simulated data: 0.918.

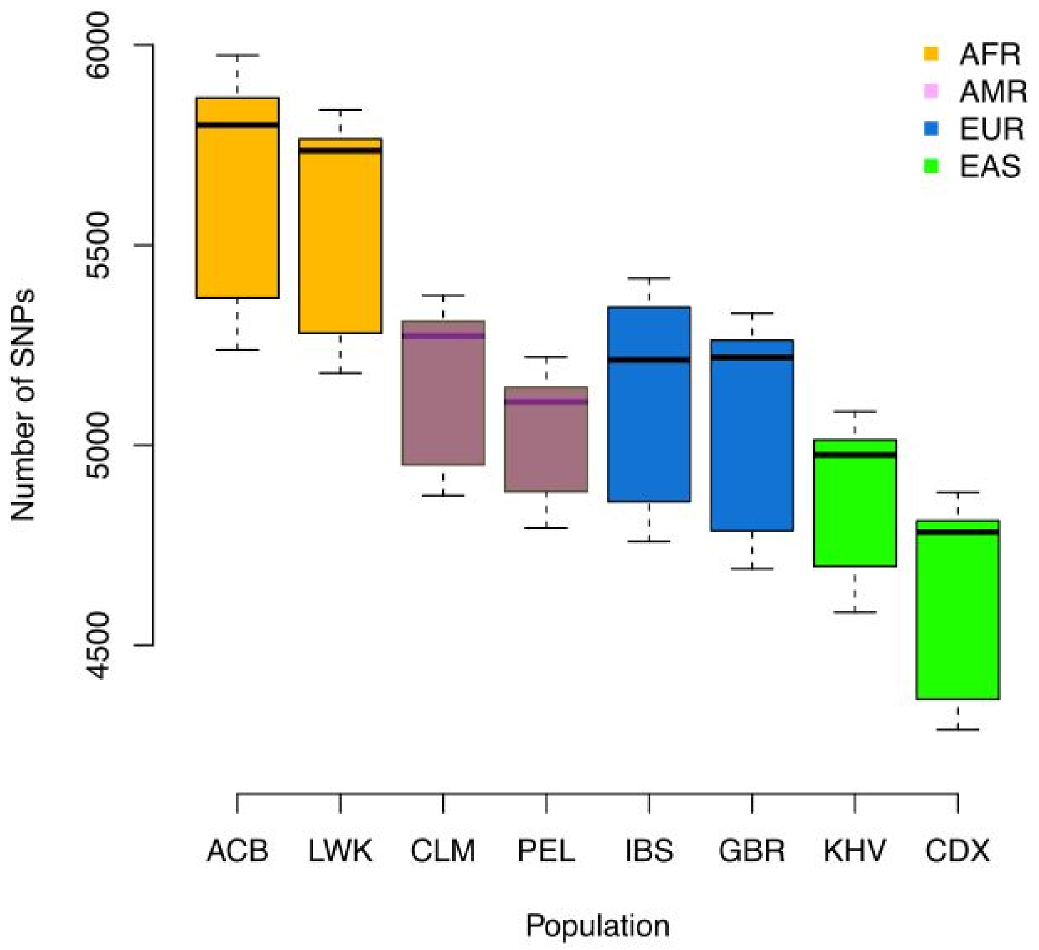


**Supplementary Figure 2**: Number of SNPs used for pairwise comparison of the simulated families per population. Each color represents a super population (AFR: African, AMR: Ad Mixed American, EUR: European, EAS: East Asian).

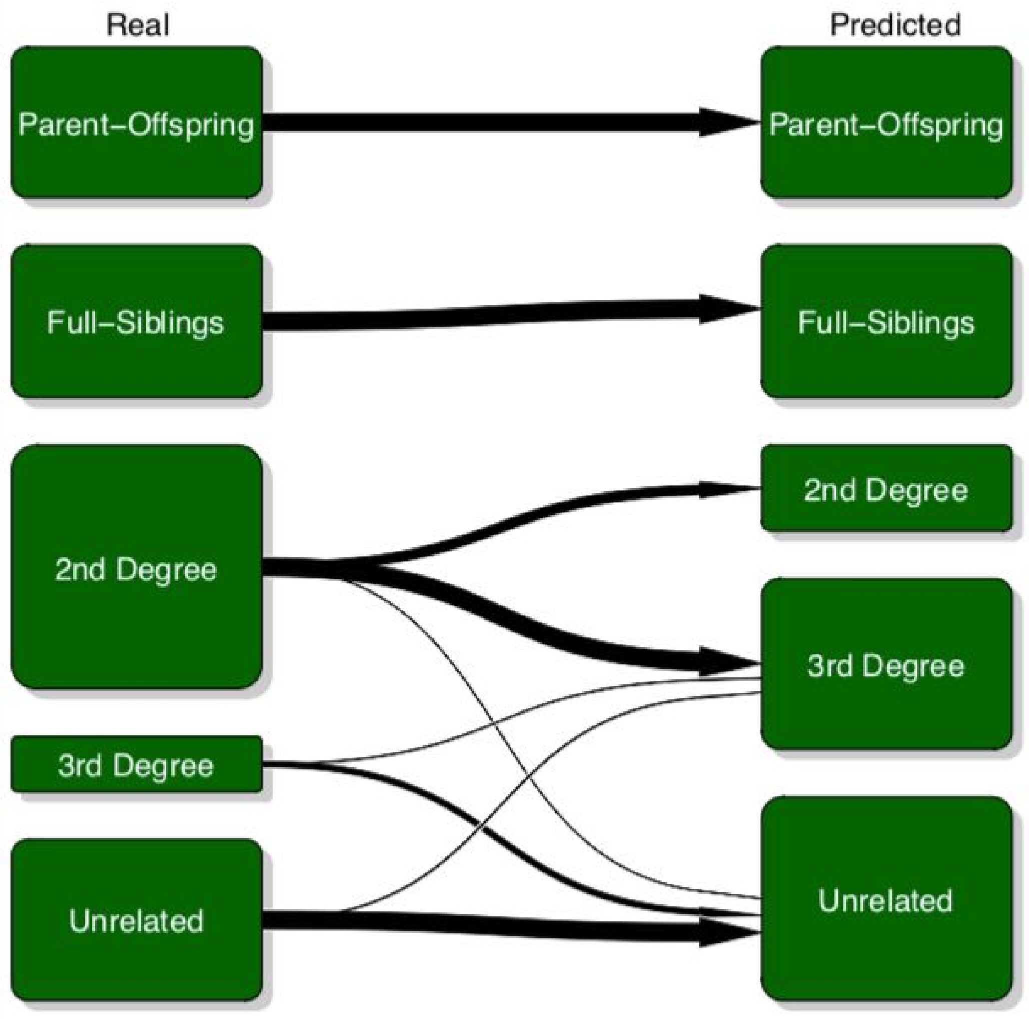


**Supplementary Figure 3**: Kinship classification by PRIMUS for simulated families (pairs with higher than third degree relationship are represented here as unrelated).
